# Identification of Hub Genes and CDK1-Targeting Therapeutics in Hepatocellular Carcinoma: Bioinformatics and Simulation Study

**DOI:** 10.1101/2024.10.10.617319

**Authors:** Ricardo Romero, Cinthia C. Toledo

## Abstract

**Background:** Hepatocellular carcinoma (HCC) remains a significant global health challenge, often associated with chronic hepatitis B (HBV) and hepatitis C (HCV) infections. Novel therapeutic approaches are urgently needed to improve patient outcomes.

**Objective:** To identify and validate potential therapeutic targets and novel drug candidates for HBV, HCV, and associated HCC using integrative bioinformatics and molecular dynamics simulations.

**Methods:** Gene expression datasets from HCC studies were analyzed to identify dysregulated genes. Protein-protein interaction networks were constructed to determine hub genes. Virtual screening of kinase compound libraries against CDK1 was performed, followed by molecular docking and dynamics simulations. Drug-likeness and metabolic properties of hit compounds were evaluated.

**Results:** Seven hub genes were identified, including five upregulated (CDK1, AURKB, CCNA2, BUB1B, RRM2) and two downregulated (AGXT2, ESR1) genes. Virtual screening yielded three promising hit compounds (G213-0272, E130-0174, Z666960592) targeting CDK1. E130-0174 demonstrated the most favorable profile, with stable CDK1 binding and no inhibition of major cytochrome P450 enzymes. Two repurposed drugs, alvocidib and riviciclib, also showed potential as CDK1 inhibitors.

**Conclusions:** This study identified novel CDK1 inhibitors with promising profiles for HCC treatment. The integrative approach combining bioinformatics and molecular dynamics simulations provides a robust framework for identifying potential therapeutic targets and drug candidates in HCC and related liver diseases. Further experimental validation is warranted to confirm these findings and advance the development of new HCC treatments.

## Introduction

Viral hepatitis, particularly hepatitis B (HBV) and hepatitis C (HCV), remains a significant global health challenge, affecting millions of individuals worldwide [1]. Chronic infection with these viruses is a leading cause of liver cirrhosis and hepatocellular carcinoma (HCC), the most common form of primary liver cancer [2]. Despite advances in antiviral therapies and vaccination programs, the burden of these diseases continues to grow, necessitating the development of novel therapeutic approaches [3], particularly for those at risk of progressing to HCC.

HBV, a DNA virus from the Hepadnaviridae family, and HCV, an RNA virus from the Flaviviridae family, both exhibit complex mechanisms of immune evasion and chronic infection. In the case of HBV, the persistence of covalently closed circular DNA (cccDNA) in hepatocytes enables viral replication even after suppression by nucleos(t)ide analog therapies, making complete viral eradication difficult [4]. HCV, on the other hand, can now be effectively cured in the majority of cases with direct-acting antivirals (DAAs) targeting the viral protease, polymerase, and nonstructural proteins (NS5A) [5]. Despite these advances, both HBV and HCV are associated with a high risk of developing HCC, especially in individuals with chronic infection and liver cirrhosis.

In HBV-related HCC, viral DNA integration into the host genome can drive oncogenesis by activating oncogenes and disrupting tumor suppressor genes [6]. Similarly, in HCV-related HCC, chronic inflammation and liver injury promote oncogenic mutations and hepatocyte proliferation. This complex interplay between viral factors, host immune responses, and hepatocyte dysfunction in the progression from chronic viral hepatitis to HCC presents numerous potential therapeutic targets [7]. However, identifying and validating these targets, as well as discovering effective drug candidates, that can target both viral replication and cancer progression, remains a significant challenge in the field.

Recent advancements in high-throughput experimental techniques have generated vast amounts of genomic, proteomic, and structural data related to HBV, HCV, and HCC [8]. Integrative bioinformatics approaches and molecular dynamics (MD) simulations offer powerful tools to analyze these data and provide insights into potential therapeutic targets and drug interactions [9, 10].

In the current study, we employed a comprehensive bioinformatics-driven approach to identify novel therapeutic targets for HBV, HCV, and associated HCC. Through the integration of transcriptomic, proteomic, and interactomic datasets, we aimed to uncover key molecular pathways that are dysregulated in these diseases. By incorporating relevant hepatitis and HCC data from publicly available studies, we identified critical hubs in the viral replication and oncogenesis networks. These hubs represent promising therapeutic targets, as they are involved in essential viral-host interactions and cancer progression mechanisms.

Our bioinformatics analysis was complemented by molecular dynamics (MD) simulations, a computational approach that allows the prediction of the structural and dynamic properties of molecular systems at the atomic level. MD simulations were used to model the binding interactions between candidate drug molecules and the identified targets, providing insights into the potential efficacy of these compounds in inhibiting viral replication and cancer-related signaling pathways. This integrative approach enabled us to prioritize therapeutic targets based on both their biological significance and druggability, paving the way for the rational design of novel therapeutics.

The primary objective of this research was to identify and validate potential therapeutic targets for HBV, HCV, and related HCC using a combination of bioinformatics analyses and MD simulations. Specifically, we aimed to:

- Identify key dysregulated genes and proteins associated with HBV, HCV, and HCC progression through integrative bioinformatic analyses.
- Analyze the molecular interactions between identified targets and candidate therapeutic compounds using molecular docking and MD simulations.
- Evaluate the stability and binding affinity of these interactions to prioritize promising drug candidates.

By integrating omics data with structural biology techniques, this study provides a comprehensive framework for the discovery of novel therapeutic targets and drug candidates. The results presented here may contribute to the development of more effective treatment strategies for viral hepatitis and HCC, particularly for patients who are unresponsive to current therapies or at high risk of cancer progression.

## Methods

### Data Acquisition and Differential Gene Expression Analysis

Publicly available gene expression datasets related to hepatitis and hepatocellular carcinoma were retrieved from the Gene Expression Omnibus (GEO) database. Specifically, studies GSE121248 [11], GSE83148 [12,13], GSE38941 [14], and GSE41804 [15] were selected for analysis. Differential gene expression analyses were conducted using the NCBI’s GEO2R online tool, applying the criteria of |Log2FC| > 0 and adjusted p-value (FDR) < 0.05. Genes showing significant differential expression were classified into upregulated and downregulated groups. Common dysregulated genes across the selected studies were identified using Funrich [16].

### Protein-Protein Interaction (PPI) Network Construction

PPI networks for upregulated and downregulated genes were constructed using the STRING database [17,18], with a high interaction confidence score of 0.7. The resulting networks were used to identify interactions between the dysregulated genes and other proteins.

### Hub Genes Identification

Hub genes were identified through Cytoscape [19] using the CytoHubba plugin [20]. Two metrics, interaction degree and betweenness centrality, were employed to rank the hub genes based on their importance in the PPI networks.

Identified hub genes were validated against data from the GEPIA2 platform [21], which incorporates gene expression data from The Cancer Genome Atlas (TCGA) and GTEx databases [22,23], as well as the Hepatocellular Carcinoma Database (HCCDB) [24], which integrates mRNA expression data from various GEO studies. Expression levels and survival analyses were performed to confirm the relevance of the hub genes in hepatocellular carcinoma.

### Target Selection and Virtual Screening

CDK1, identified as a top hub gene, was selected as a target for virtual screening. The crystal structure of CDK1 (PDB ID: 4y72 [25]) was retrieved from the Protein Data Bank and prepared for docking. The structure was cleaned of solvents and ligands, energy minimized by the steepest descent method using the GROMOS force field [26] via the SwissPDB tool [27], and processed for docking using UCSF Chimera [28].

### Compound Library Preparation and Filtering

Kinase compound libraries were downloaded from ChemDiv [29] and Enamine [30]. The compounds were first screened for toxicity properties such as tumorigenic, mutagenic, irritant, and reproductive effects using the DataWarrior software [31]. Subsequently, drug-likeness features were assessed based on Lipinski’s Rule of Five [32], Ghose’s [33], Veber’s [34], Egan’s [35], and Muegge’s rules [36]. Compounds were also filtered for PAINS (Pan Assay Interference Compounds) [37], Brenk’s structural alerts [38], and lead-likeness, using a combination of Python scripts and the SwissADME server [39]. We also included five drugs targeting CDK1 [40], Avotaciclib, Alvocidib, Roniciclib, Riviciclib, and Dinaciclib, both as controls and for repurposed analyses. Ligands were energy minimized using the Generalized Amber Force Field (GAFF) by the steepest descent method, and prepared for docking with hydrogens and charges added via the Open Babel software [41].

### Deep Learning-Based Bioactivity Prediction

In addition to the filtering process, compounds were subjected to a bioactivity assessment using a deep learning model [42], trained on tyrosine kinase compounds [43], to evaluate their potential activity against CDK1.

### Molecular Docking and Dynamics Simulations

Docking simulations were performed using AutoDock Vina [44,45] to predict the binding affinity of the selected compounds to the CDK1 active site. The top-scoring compounds were further analyzed through molecular dynamics (MD) simulations using the GROMACS software [46] to evaluate the stability and behavior of the ligand-protein complexes in a dynamic environment.

## Results

### Differential Gene Expression Analysis

Analysis of the four GEO studies revealed a total of 1,757 genes that were consistently dysregulated across all datasets. Of these, 852 genes were upregulated, and 905 genes were downregulated. These genes formed the basis for subsequent protein-protein interaction (PPI) network analysis. (Figure 2).

**Figure 1.**
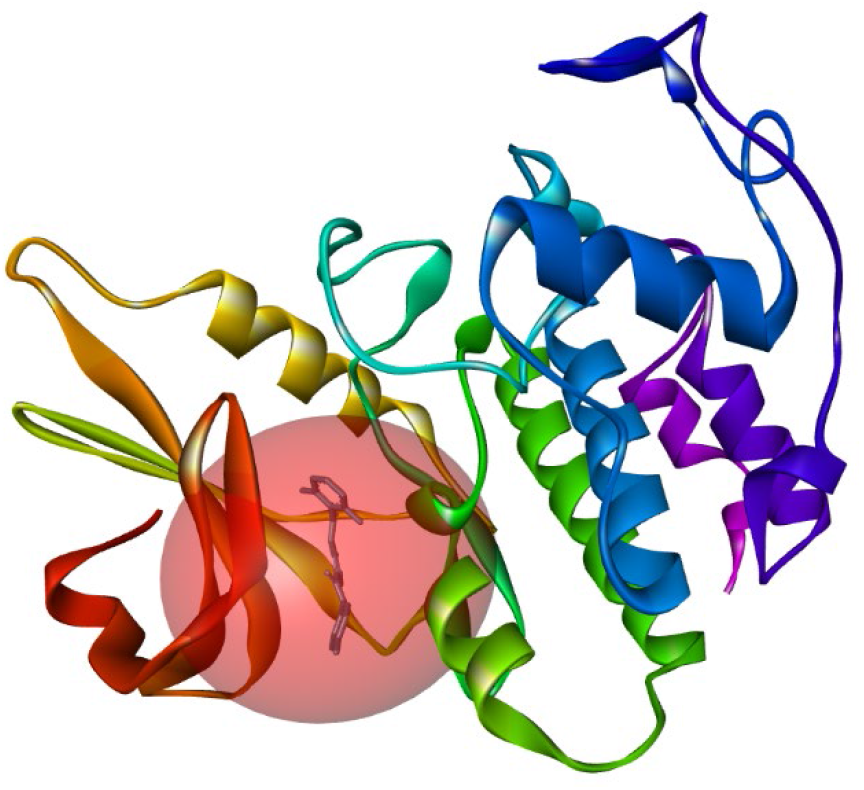
Target structure. Crystallized structure of CDK1 in complex with a ligand (PDB 4Y72), showing the protein’s active site.

**Figure 2.**
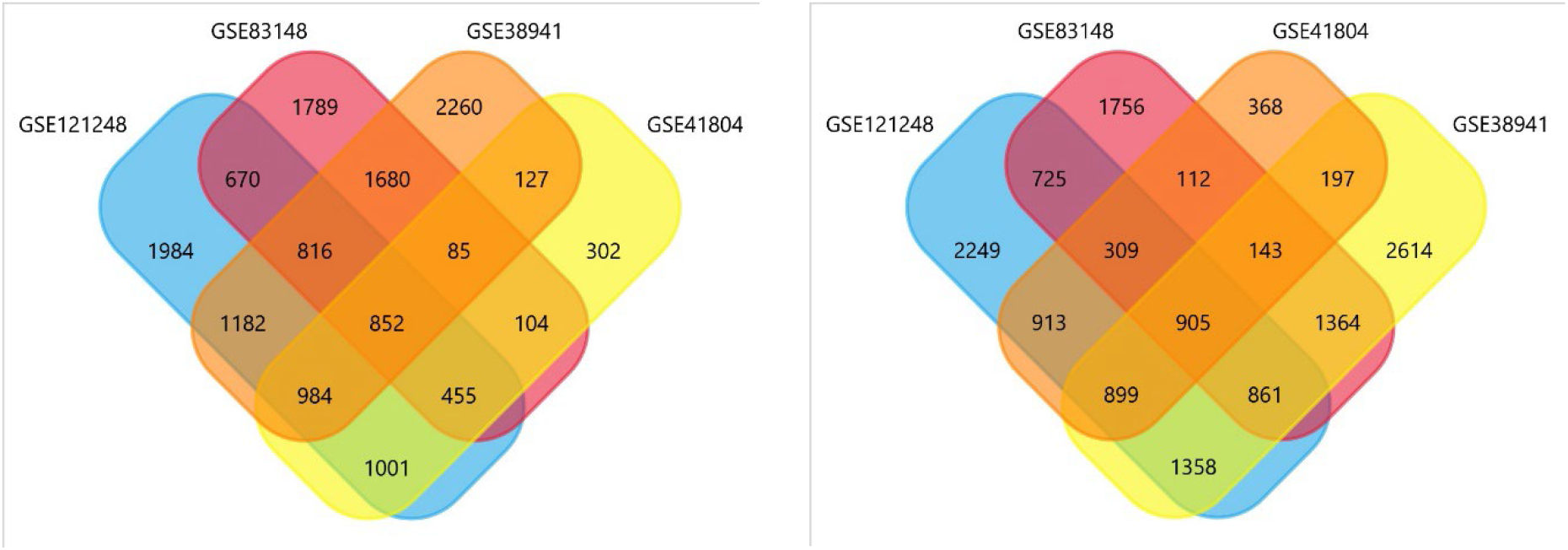
Venn diagrams of dysregulated genes. The image on the left corresponds to upregulated genes in hepatitis and hepatocellular carcinoma (HCC) from GEO studies (GSE121248, GSE83148, GSE38941, and GSE41804). The overlapping areas indicate common genes across the datasets. The image on the right is for downregulated genes in hepatitis and HCC from the same GEO studies. The overlaps similarly represent common downregulated genes shared among the datasets. These diagrams help visualize the shared and unique gene expression patterns across different studies in the context of hepatitis and HCC.

### Hub Genes Identification

PPI network analysis of the dysregulated genes yielded significantly enriched networks (provided as supplementary material). From these networks, we identified seven hub genes:

Upregulated hub genes:

- CDK1 (Cyclin Dependent Kinase 1)
- AURKB (Aurora Kinase B)
- CCNA2 (Cyclin A2)
- BUB1B (BUB1 Mitotic Checkpoint Serine/Threonine Kinase B)
- RRM2 (Ribonucleotide Reductase Regulatory Subunit M2)

Downregulated hub genes:

- AGXT2 (Alanine-Glyoxylate Aminotransferase 2)
- ESR1 (Estrogen Receptor 1)

The PPI network of the upregulated hub genes demonstrated a highly interconnected cluster, suggesting potential functional relationships or co-regulation among these genes in HCC (Figure 3).

**Figure 3.**
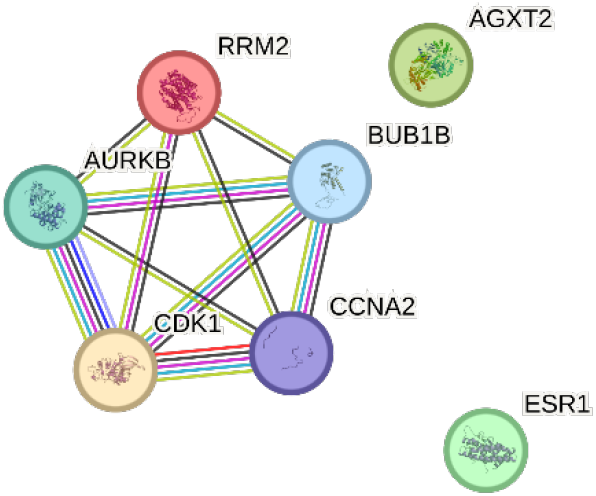
Hub genes PPI network. This figure represents the Protein-Protein Interaction (PPI) network of HCC hub genes. The genes that are connected in the network (CDK1, CCNA2, AURKB, BUB1B, RRM2) are upregulated in hepatocellular carcinoma (HCC), forming a highly interconnected cluster, with an interaction score of 0.7 and a PPI enrichment p value of 1.37e-08. The disconnected genes (AGXT2 and ESR1) are downregulated and do not show direct interactions with the upregulated genes in this PPI network. This network highlights the functional relationships among the upregulated hub genes involved in cell cycle regulation and proliferation in HCC.

### Survival Analysis

Independent validation of the hub genes from HCCDB confirmed their dysregulation in HCC. The upregulated hub genes (CDK1, AURKB, CCNA2, BUB1B, and RRM2) showed significantly higher expression levels in HCC tissues compared to normal liver tissues (Figure 4).

**Figure 4.**
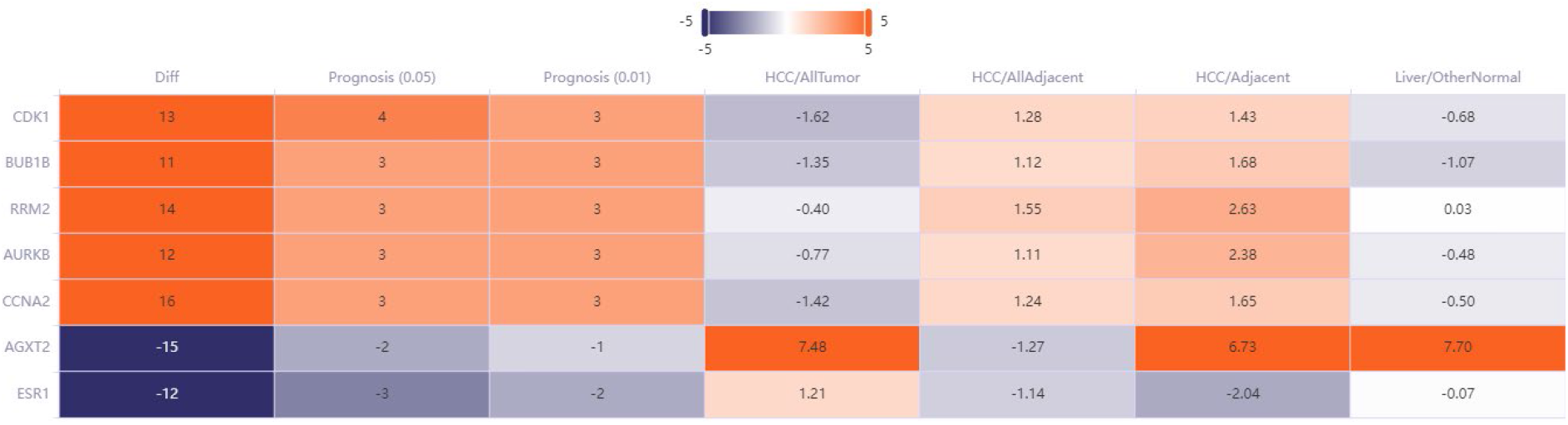
Expression heatmap. The heatmap illustrates the expression profiles of HCC hub genes across various conditions and prognostic categories. The hub genes include CDK1, BUB1B, RRM2, AURKB, CCNA2, AGXT2, and ESR1. The columns represent the following categories: **Diff**: Differential expression in HCC versus normal tissue. **Prognosis (0.05) & Prognosis (0.01)**: Association with prognosis at significance thresholds of 0.05 and 0.01. **HCC/AllTumor**: Expression in HCC tumor samples. **HCC/AllAdjacent**: Expression in tissue adjacent to the tumor. **HCC/Adjacent**: Expression in adjacent non-tumor tissue. **Liver/OtherNormal**: Expression in normal liver and other non-cancerous tissues. Positive values in red represent upregulation, while negative values in blue represent downregulation. CDK1, BUB1B, RRM2, AURKB, and CCNA2 are generally upregulated in tumor tissues, whereas AGXT2 shows a marked increase in non-tumor liver tissues.

Survival analysis using the GEPIA2 platform confirmed that the upregulated hub genes were linked to poorer prognosis in HCC patients, while the downregulated genes were associated with better outcomes. Specifically, CDK1, BUB1B, RRM2, AURKB, and CCNA2 had positive hazard ratios (HRs), indicating that higher expression levels correlated with worse survival, whereas AGXT2 and ESR1 had negative HRs, suggesting a protective role (Figure 5).

**Figure 5.**
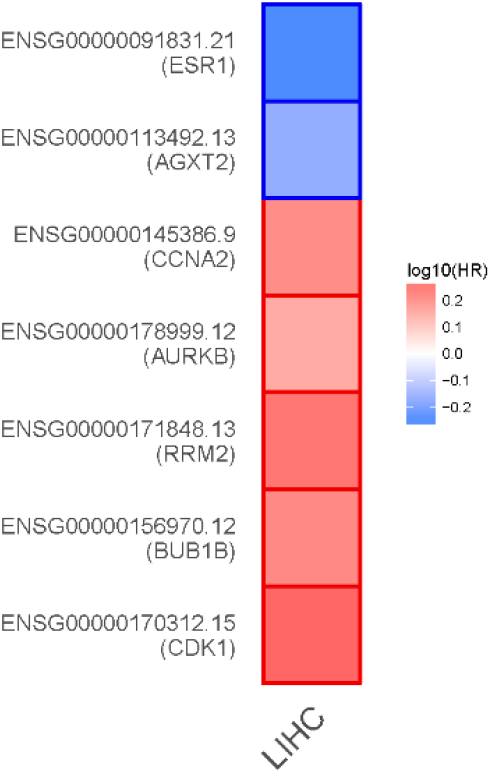
Survival heat map. This map represents the log10 hazard ratios (HR) for hub genes in liver hepatocellular carcinoma (LIHC). The genes shown include ESR1, AGXT2, CCNA2, AURKB, RRM2, BUB1B, and CDK1. The color gradient represents the hazard ratio: Red shades indicate genes associated with increased risk (positive HR) and poorer survival outcomes. Blue shades indicate genes associated with decreased risk (negative HR) and better survival outcomes. Notably, CDK1, BUB1B, RRM2, AURKB, and CCNA2 are all linked with higher hazard ratios (positive, in red), suggesting they are associated with poorer prognosis in LIHC. In contrast, ESR1 and AGXT2 show negative hazard ratios (in blue), implying better survival outcomes for patients with higher expression of these genes.

### Virtual Screening Results

Following toxicity screening, drug-likeness filtering, and molecular docking of the ChemDiv and Enamine libraries against the CDK1 protein structure yielded three promising hit compounds: G213-0272 (ChemDiv library), E130-0174 (ChemDiv library) and Z666960592 (Enamine library). Table I provides the binding energy to CDK1 from molecular docking of the identified hits and the control/repurposed drugs.

**Table I.** Compounds IDs. Names, SMILES and docking binding energy to CDK1, in kcal/mol, of identified hits and control / repurposed drugs. For the hits, the compound name refers to the chemical library ID.

| Compound | SMILES | Binding Energy |
| --- | --- | --- |
| **G213-0272** | CN1CCN(CCNC(c2noc-3c2CCc2c3ccs2)=O)CC1 | -8.49 |
| **E130-0174** | Cc(cc1)cnc1NC(CCc1nc(C2=CCN(C)CC2)no1)=O | -9.27 |
| **Z666960592** | NC(C(CC1)CN1C(c1cc2c(-c3ccccc3)[o]nc2cc1)=O)=O | -10.90 |
| **Avotaciclib** | C1=CC(=NC(=C1O)C2=NC(=NC=C2)N)C3=NC(=NC=C3)N | -9.09 |
| **Alvocidib** | CN1CC[C@@H]([C@@H](C1)O)C2=C(C=C(C3=C2OC(=CC3=O)C4=CC=CC=C4Cl)O)O | -11.26 |
| **Roniciclib** | C[C@H]([C@@H](C)OC1=NC(=NC=C1C(F)(F)F)NC2=CC=C(C=C2)S(=N)(=O)C3CC3)O | -9.76 |
| **Rivaciclib** | CN1CC[C@H]([C@@H]1CO)C2=C(C=C(C3=C2OC(=CC3=O)C4=CC=CC=C4Cl)O)O | -8.93 |
| **Dinaciclib** | CCC1=C2N=C(C=C(N2N=C1)NCC3=C[N+](=CC=C3)[O-])N4CCCC[C@H]4CCO | -8.30 |

Ligand 2 (E130-0174) demonstrated the lowest fluctuations in Root Mean Square Deviation (RMSD), indicating strong stability within the CDK1 binding site, while Ligand 3 (Z666960592) showed higher structural deviations (Figure 6). The potential energy analysis further supported that Ligand 2 had the most stable interaction with CDK1, suggesting stronger binding affinity compared to the other compounds (Figure 7).

**Figure 6.**
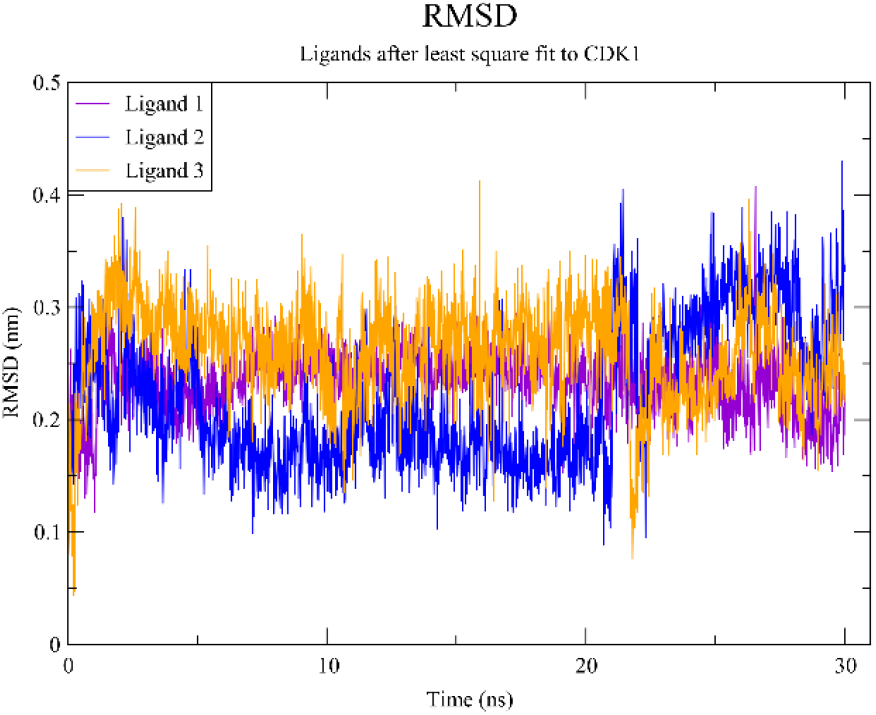
Root Mean Square Deviation (RMSD). The plot shows the RMSD of the three hits (Ligand 1, Ligand 2, and Ligand 3, in the same order of Table I) after least-square fitting to the CDK1 protein. The plot shows the structural stability of the ligands over a 30 ns molecular dynamics simulation. Ligand 2 (blue) shows the least fluctuation, indicating higher stability, while Ligand 3 (orange) exhibits the highest fluctuations in RMSD, indicating greater structural deviations. RMSD values range from 0.1 nm to 0.4 nm, reflecting how each ligand behaves within the protein’s binding pocket over time.

**Figure 7.**
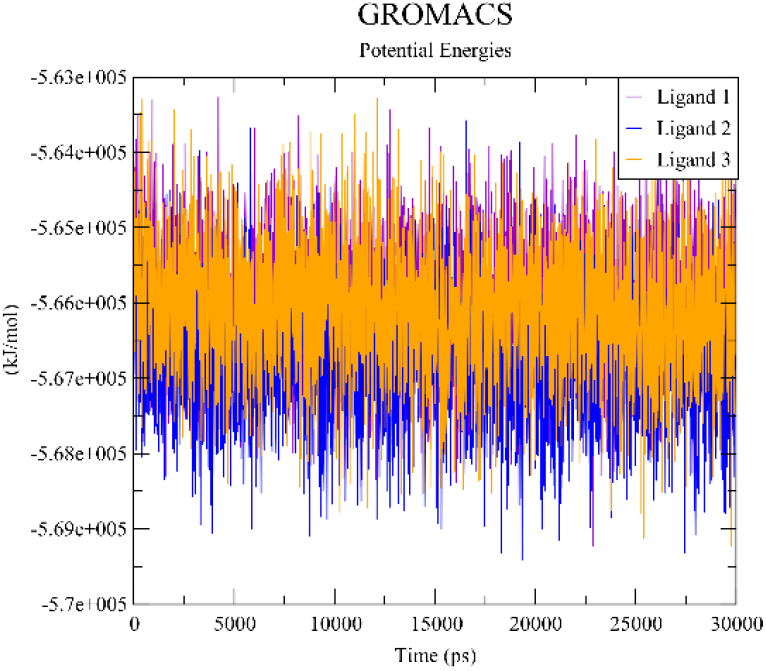
Potential energy plot. The plot shows the potential energy of the three hits (Ligand 1, Ligand 2, and Ligand 3, in the same order of Table I) interacting with CDK1 during a 30 ns molecular dynamics simulation. Ligand 2 (blue) exhibits the lowest and most stable potential energy values, suggesting stronger binding affinity and stability with the protein. Ligand 3 (orange) has higher and more fluctuating potential energy, indicating less stable binding. Ligand 1 (purple) remains between the other two ligands, showing moderate stability throughout the simulation.

All three lead compounds passed toxicity assessments, showing no tumorigenic, mutagenic, irritant, or reproductive effects. They also satisfied drug-likeness criteria based on Lipinski’s Rule of Five and related guidelines. Compound E130-0174 exhibited the best overall drug-likeness and solubility profile (Table II).

**Table II.** Toxicity and Druglikeness properties. The toxicity column includes results for mutagenic, tumorigenic, irritant and reproductive effectiveness. Toxicity, Solubility, Druglikeness and Drug-score values were obtained with the Osiris Property Explorer software. Avotaciclib exhibited one violation of Egan’s rules, and also Roniciclib, plus one Ghose violation. Dinaciclib exhibited two Brenk alerts. All the compounds were classified as active by the deep learning bioactivity model.

| Compound | Toxicity | Solubility | Druglikeness | Drug-score | MW | TPSA |
| --- | --- | --- | --- | --- | --- | --- |
| **G213-0272** | None | -2.07 | 10.72 | 0.9 | 346.45 | 89.85 |
| **E130-0174** | None | -1.36 | 5.98 | 0.92 | 327.38 | 84.15 |
| **Z666960592** | None | -4.03 | 7.71 | 0.79 | 335.36 | 89.43 |
| **Avotaciclib** | None | -2.15 | -1.77 | 0.54 | 281.27 | 136.72 |
| **Alvocidib** | None | -3.87 | 6.97 | 0.7 | 401.84 | 94.14 |
| **Roniciclib** | None | -5.46 | -6.23 | 0.27 | 430.44 | 116.57 |
| **Rivaciclib** | None | -3.87 | 6 | 0.71 | 401.84 | 94.14 |
| **Dinaciclib** | Reproductive | -7.1 | 0.16 | 0.3 | 396.49 | 91.15 |

In addition to the novel compounds, two control/repurposed drugs, Alvocidib and Riviciclib also passed the established filters and showed favorable results in molecular dynamics simulations (provided as supplementary material), suggesting their potential efficacy as CDK1 inhibitors in HCC treatment.

### Metabolic Properties and Cytochrome P450 Interaction

The metabolic properties of the lead compounds and the repurposed drugs were evaluated with respect to their inhibition of cytochrome P450 (CYP) enzymes. G213-0272 was found to inhibit both CYP1A2 and CYP3A4, while E130-0174 showed no inhibition of major CYP enzymes, making it a safer candidate in terms of potential drug-drug interactions (Table III).

**Table III.**
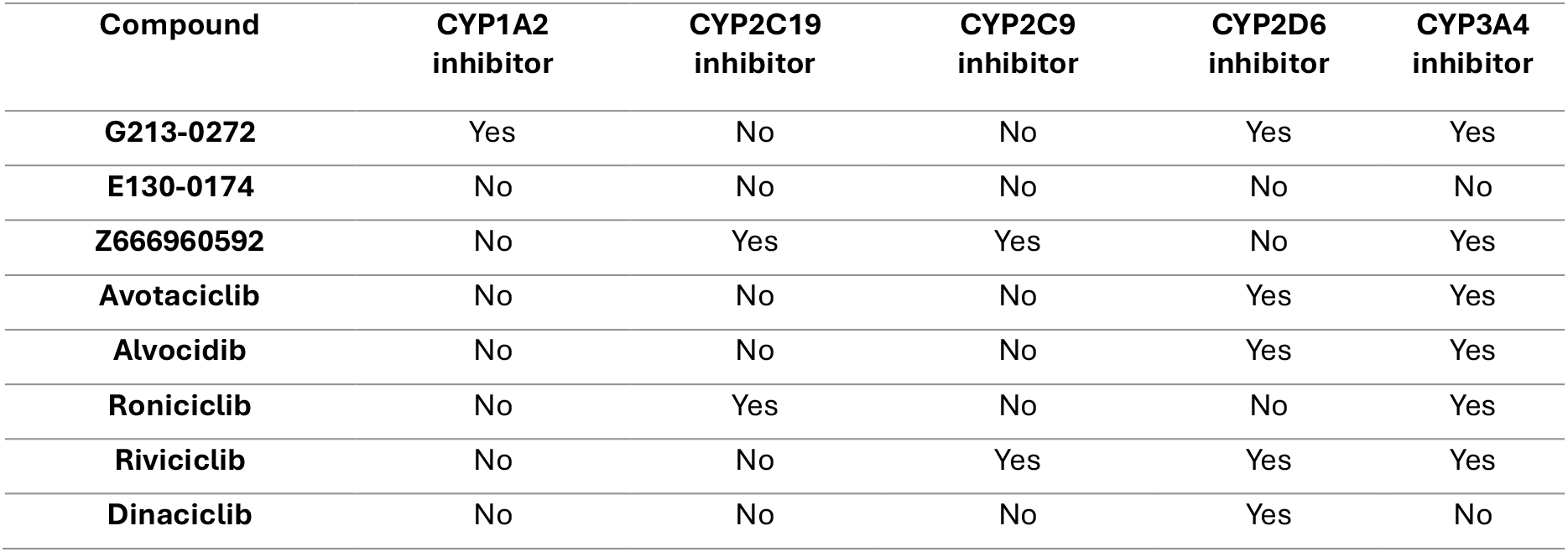
Metabolic Properties. Hits and controls metabolic properties regarding cytochrome p450. The columns indicate whether or not the compound inhibits the corresponding enzyme.

| Compound | CYP1A2 inhibitor | CYP2C19 inhibitor | CYP2C9 inhibitor | CYP2D6 inhibitor | CYP3A4 inhibitor |
| --- | --- | --- | --- | --- | --- |
| **G213-0272** | Yes | No | No | Yes | Yes |
| **E130-0174** | No | No | No | No | No |
| **Z666960592** | No | Yes | Yes | No | Yes |
| **Avotaciclib** | No | No | No | Yes | Yes |
| **Alvocidib** | No | No | No | Yes | Yes |
| **Roniciclib** | No | Yes | No | No | Yes |
| **Rivaciclib** | No | No | Yes | Yes | Yes |
| **Dinaciclib** | No | No | No | Yes | No |

Overall, the integrative bioinformatics and molecular simulation approach allowed us to identify novel therapeutic targets and prioritize drug candidates with promising efficacy for the treatment of HCC.

## Discussion

### Identified hub genes

This study employed an integrative bioinformatics and molecular dynamics approach to identify potential therapeutic targets and novel drug candidates for hepatitis B virus (HBV), hepatitis C virus (HCV), and associated hepatocellular carcinoma (HCC). Our analysis revealed several upregulated hub genes, including CDK1, AURKB, CCNA2, BUB1B, and RRM2, underscores their potential as therapeutic targets. These genes were found to be highly interconnected within the protein-protein interaction (PPI) network, indicating their potential functional relationships or co-regulation in HCC pathogenesis.

The overexpression of these hub genes correlates with poorer survival outcomes in HCC, as confirmed through independent validation with the GEPIA2 platform and survival analysis (Figure 5). For example, CDK1 is a critical regulator of mitosis [47], and its inhibition could lead to cell cycle arrest, making it a compelling target for therapeutic intervention in HCC. AURKB, involved in chromosome segregation and cytokinesis, has been associated with early HCC recurrence [48]. The cyclin A2 gene (CCNA2) plays a crucial role in cell cycle regulation and has been identified as a prognostic marker in HCC [49]. The integration of these findings into clinical practice could involve developing inhibitors against these upregulated genes or leveraging them for diagnostic purposes to predict disease progression and treatment responses.

For the downregulated hub genes AGXT2 and ESR1, their lower expression levels were associated with improved survival outcomes in HCC patients. These genes could serve as valuable biomarkers for early-stage HCC or for identifying patients who may benefit from less aggressive treatment strategies. AGXT2 has been implicated in metabolic reprogramming in HCC, and its downregulation may serve as an early marker of hepatocarcinogenesis [50]. ESR1, encoding estrogen receptor α, has been associated with gender disparity in HCC incidence and could be explored as a therapeutic target [51]. Biomarker-driven treatment strategies have gained prominence, and incorporating these downregulated genes into clinical screening tools could enhance early detection and personalized treatment options [52].

### Identified hits

The virtual screening and molecular dynamics simulations identified several promising compounds targeting CDK1, with E130-0174 emerging as a standout hit due to its strong binding affinity and stable interaction within the CDK1 binding pocket. Importantly, E130-0174 does not inhibit any of the major cytochrome P450 enzymes (CYP1A2, CYP2C19, CYP2C9, CYP2D6, or CYP3A4), making it a safer candidate with reduced risk of drug-drug interactions. This is particularly important in HCC treatment, where patients often require multiple medications and have compromised liver function [53]. In contrast, G213-0272 inhibits CYP1A2, CYP2D6, and CYP3A4, while Z666960592 inhibits CYP2C19, CYP2C9, and CYP3A4, which could lead to interactions with other drugs metabolized by these enzymes. Among the repurposed drugs, Alvocidib and Riviciclib show strong binding to CDK1, but also inhibit CYP2D6 and CYP3A4, increasing the risk of drug interactions in polypharmacy contexts often seen in HCC treatment.

Additionally, the molecular dynamics simulations reinforced the stability of the ligand-protein complexes. E130-0174 demonstrated the most stable interaction with CDK1, with minimal fluctuations in Root Mean Square Deviation (RMSD) and the lowest potential energy values, suggesting stronger binding affinity. These characteristics make E130-0174 a promising lead compound for further development. The identification of these CDK1 inhibitors is significant not only for HCC but also for other cancers where CDK1 plays a crucial role in cell cycle progression and survival [40,54]. On the other hand, control drugs like Dinaciclib and Roniciclib, while potent CDK1 inhibitors, presented poor druglikeness scores and some violations of druglikeness filters, limiting their potential as viable therapeutic options.

### Clinical implementations

While the bioinformatics-driven identification of CDK1 as a hub gene and therapeutic target is a valuable contribution to hepatocellular carcinoma (HCC) research, integrating these findings into clinical practice requires a more in-depth exploration of their applicability. One potential avenue for immediate clinical relevance lies in using CDK1 and other identified hub genes (e.g., AURKB, RRM2) as biomarkers for patient stratification. Biomarker-driven approaches have transformed the management of several cancers, and this is particularly urgent for HCC, which currently lacks molecular biomarkers for personalized treatment [53]. For instance, patients with high CDK1 expression could be more likely to respond to CDK1-targeted therapies, while those with lower expression might benefit from alternative treatment strategies. Future studies should focus on validating these hub genes as predictive biomarkers in clinical HCC samples, correlating their expression levels with patient outcomes such as response to existing therapies, progression-free survival, and overall survival.

In addition to the identification of potential biomarkers, there is significant promise in exploring combination therapies that target CDK1 alongside established treatments for HCC. For example, CDK1 inhibition could be tested in combination with sorafenib, which is a standard first-line treatment for advanced HCC. Preclinical studies have already shown that CDK1 inhibition can sensitize cancer cells to chemotherapy and other treatments by impairing cell cycle progression and promoting apoptosis [40]. Such a combination strategy could enhance therapeutic efficacy, particularly in patients who develop resistance to current therapies. Similarly, combining CDK1 inhibitors with emerging immunotherapies, such as immune checkpoint inhibitors [52], could exploit the tumor’s impaired cell cycle control to augment immune-mediated tumor destruction. Thus, further research into how CDK1 inhibitors can synergize with existing therapies would provide significant insights into overcoming drug resistance and improving patient outcomes in HCC.

Moreover, the role of CDK1 as a key regulator of cell division makes it a viable target not only for monotherapy but also as part of a broader treatment strategy targeting multiple cell cycle regulators. The upregulation of other hub genes like AURKB and CCNA2 indicates that the dysregulation of the cell cycle is a central feature of HCC pathogenesis. Targeting multiple points in the cell cycle regulatory network may provide a more effective means of controlling tumor growth [53]. This multi-targeted approach could also reduce the likelihood of resistance development, which is a common issue in monotherapy strategies. Clinical trials evaluating multi-agent regimens that include CDK1 inhibitors are therefore a crucial next step.

### Study limitations

The results of this study present several promising avenues for HCC treatment; however, there are limitations that should be acknowledged. First, our findings are based on computational and bioinformatics analyses of publicly available datasets, and while these provide a robust starting point, they should be complemented by experimental validation.

Another limitation is the focus on a single target (CDK1) among the identified hub genes. Future studies should explore the potential of targeting multiple hub genes simultaneously or in combination with CDK1 inhibition. This could lead to more effective treatment strategies that address the complex molecular landscape of HCC.

### Outlook

Future studies should include in vitro assays to confirm the binding affinity and specificity of the identified compounds to CDK1. Cell-based assays using HCC cell lines would be crucial to evaluate the compounds’ effects on cell proliferation, cell cycle progression, and apoptosis. Additionally, testing these compounds in combination with current standard-of-care treatments for HCC, such as sorafenib or immunotherapies, could reveal potential synergistic effects.

In vivo studies using xenograft models or genetically engineered mouse models of HCC would be essential to assess the efficacy and toxicity profiles of these compounds. Such studies should evaluate tumor growth inhibition, survival outcomes, and potential side effects. Furthermore, pharmacokinetic and pharmacodynamic studies in animal models would provide crucial information on the compounds’ behavior in a physiological context.

Finally, integrating survival analysis with the molecular data presented in this study is an essential step toward validating these findings in clinical practice. The survival analysis included in the paper, which links high CDK1 expression to poorer prognosis, underscores the potential of this gene as a prognostic biomarker. However, additional analyses incorporating larger patient cohorts and more detailed clinical variables, such as stage at diagnosis and treatment history, would provide more robust evidence of the prognostic significance of CDK1 [54]. Future studies could also explore the relationship between CDK1 expression and patient response to specific treatments, offering a pathway toward using CDK1 as a predictive marker for therapy selection. Such work could pave the way for personalized treatment protocols, where CDK1 expression guides the choice of therapy in HCC.

## Conclusion

The integrative approach employed in this study, combining bioinformatics, molecular docking, and molecular dynamics simulations, provides a powerful framework for identifying novel therapeutic targets and potential drug candidates. The identification of CDK1 as a key hub gene and the discovery of promising inhibitors such as E130-0174 provide a strong foundation for further preclinical and potentially clinical investigations, paving the way for new therapeutic strategies in the management of HCC and related liver diseases, and also for other cancers where CDK1 plays a central role.

## Supporting information

Supplemental figures and plots

## Authors declare no conflict of interest

Ethical approval was not required since no animals or humans were used as subjects in the present study.

## Authors’ contribution

Ricardo Romero

Conceptualization (Lead), Data curation (Equal), Formal analysis (Equal), Investigation (Equal), Methodology (Equal), Project administration (Lead), Supervision (Lead), Validation (Equal), Writing – original draft (Lead), Writing – review & editing (Lead).

Cinthia C. Toledo

Conceptualization (Supporting), Data curation (Equal), Formal analysis (Equal), Investigation (Equal), Methodology (Equal), Project administration (Supporting), Validation (Equal), Writing – original draft (Supporting), Writing – review & editing (Supporting).

