## Supplemental figures and plots for "Identification of Hub Genes and CDK1-Targeting Therapeutics in Hepatocellular Carcinoma: Bioinformatics and Simulation Study"

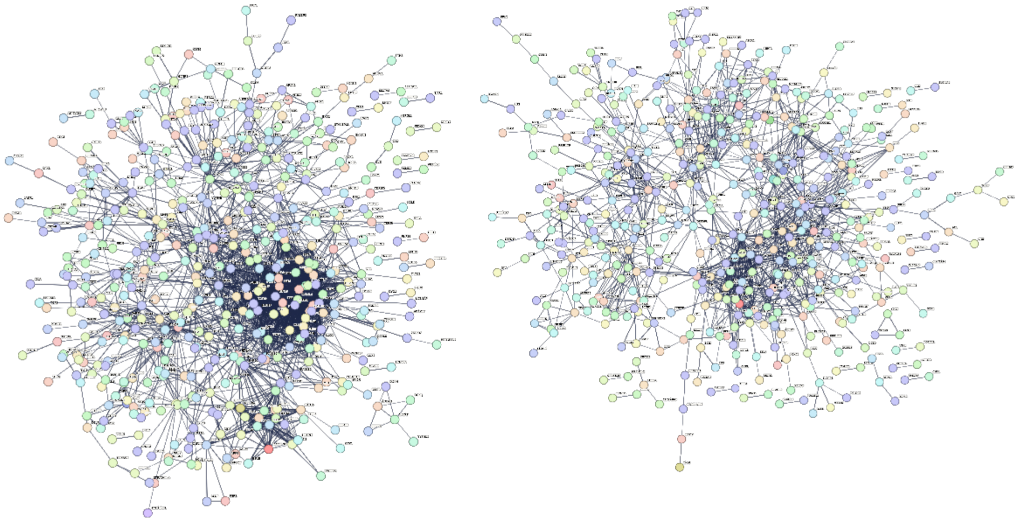

Figure 1S. PPI network of upregulated genes (left) and downregulated ones (right) common to the four GEO studies GSE121248, GSE83148, GSE3894, and GSE41804. The networks were obtained in StringDB with a high confidence score of 0.7. The upregulated network has 790 nodes and 2464 edges, with a PPI enrichment p value less than 1.0e-16. The downregulated network has 835 nodes and 1288 edges, with a PPI enrichment value less than 1.0e-16.

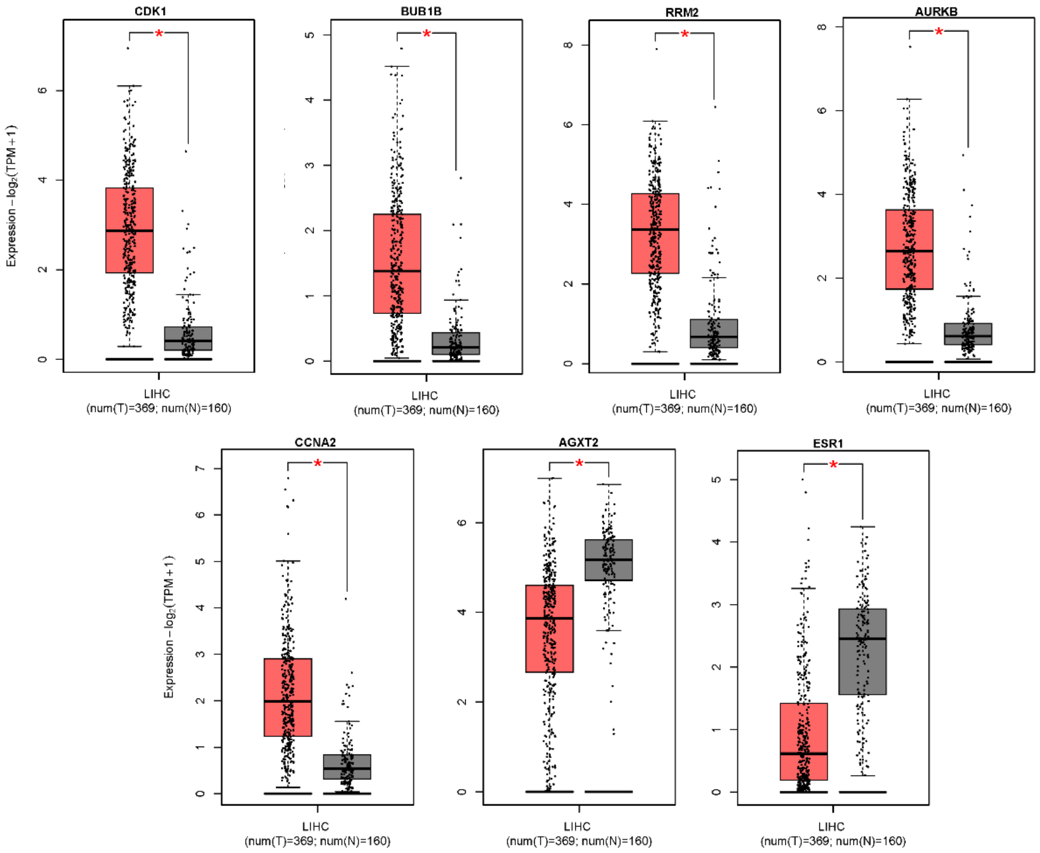


Figure 2S. Hub genes expression levels boxplots of Tumor (T in salmon) vs Normal samples (N in gray), from GEPIA2 data. The red asterisk indicates significance (adjusted p value < 0.05).

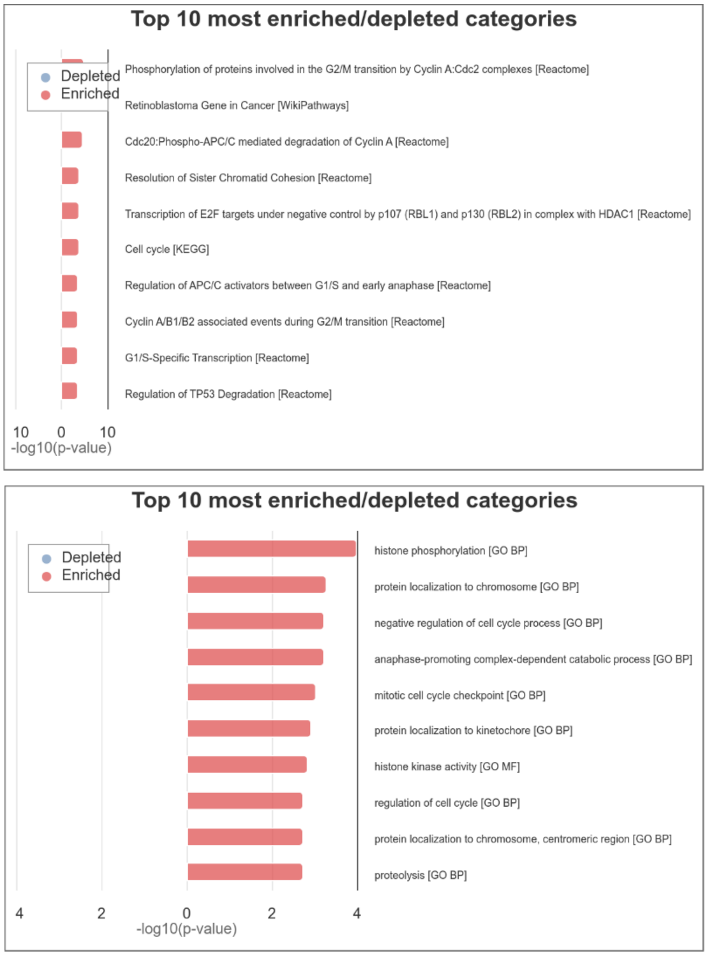


Figure 3S. Hub genes enriched categories for pathways (above) and Gene Ontology (GO, below), highlighting phosphorylation, cell cycle regulation and mitotic cell cycle. These enriched categories are central to HCC pathogenesis. The dysregulation of phosphorylation pathways, cell cycle control, and mitotic processes contributes to the hallmarks of cancer in HCC, including sustained proliferative signaling, evasion of growth suppressors, and genomic instability.

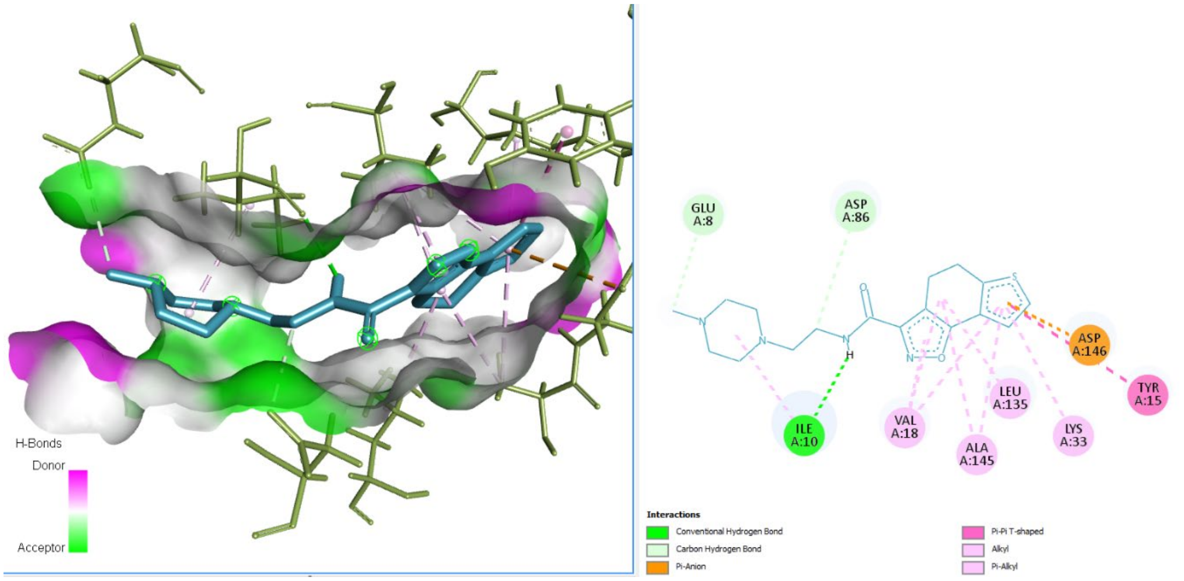


Figure 4S. Docked complex of hit G213-0272 with CDK1. On the left is the 3D view of the complex showing the surface for Hydrogen bonds. On the right there is the 2D view of interactions between the ligand and the protein. Notably, there are several Pi interactions with the aromatic rings, which contributes to the ligand's stability, further confirmed by the molecular dynamics simulation.

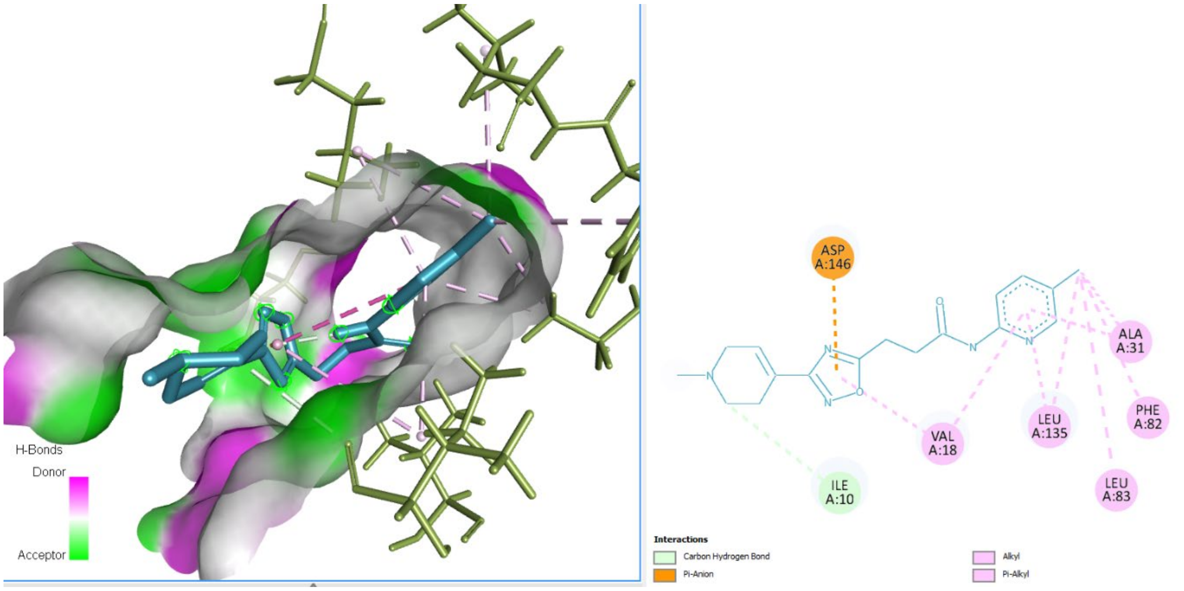


Figure 5S. Docked complex of hit E130-0174 with CDK1. On the left is the 3D view of the complex showing the surface for Hydrogen bonds. On the right there is the 2D view of interactions between the ligand and the protein. Here, there are also several Pi interactions with the aromatic rings, which contributes to the ligand's stability, further confirmed by the molecular dynamics simulation.

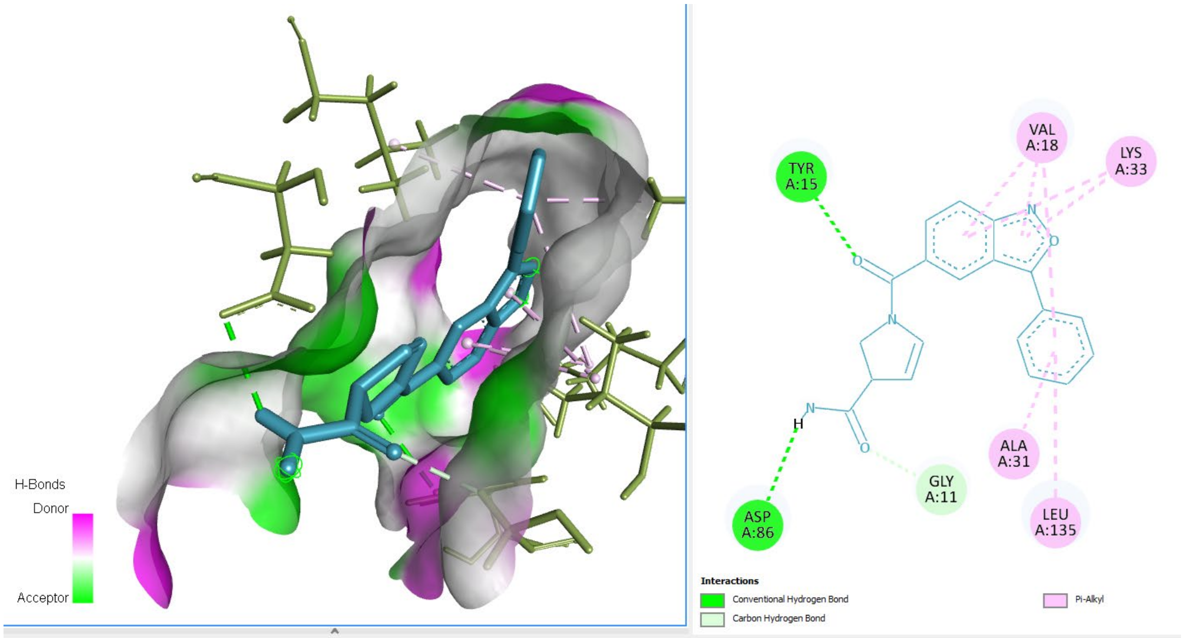


Figure 6S. Docked complex of hit Z666960592 with CDK1. On the left is the 3D view of the complex showing the surface for Hydrogen bonds. On the right there is the 2D view of interactions between the ligand and the protein. Here, there are also several Pi interactions with the aromatic rings, which contributes to the ligand's stability, further confirmed by the molecular dynamics simulation.

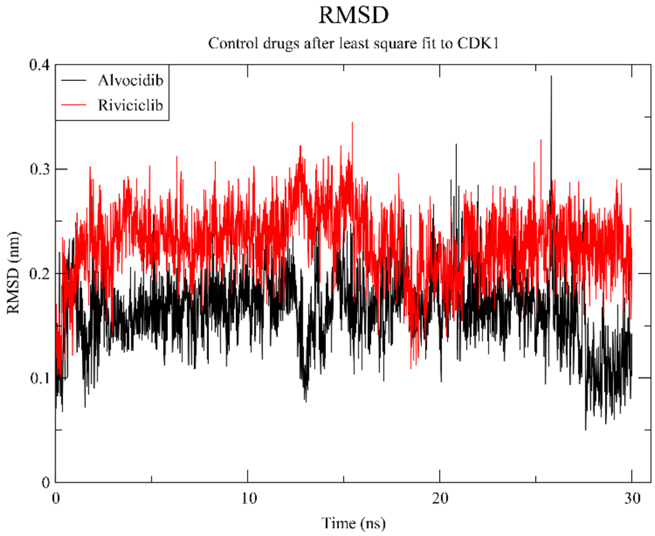


Figure 7S. Root mean square deviation of control drugs Alvocidib and Riviciclib to CDK1, during a 30 ns molecular dynamics simulation. Both drugs show great stability, with values below 0.4 nm. Alvocidib is the most stable of the two.

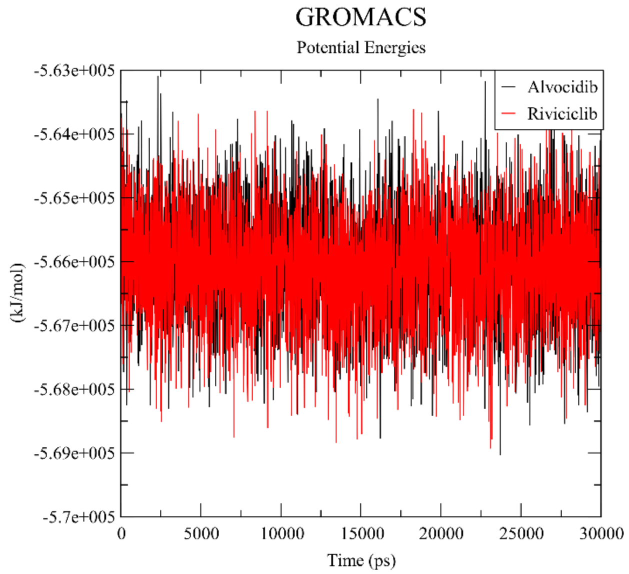


Figure 8S. Potential energies, in kJ/mol, of control drugs Alvocidib and Riviciclib bound to CDK1, during a 30 ns molecular dynamics simulation. Both drugs show great stability, with very similar binding energies of around -5.66e+005.
